# A dataset of new occurrence records of primates from the arc of deforestation, Brazil

**DOI:** 10.1101/2023.06.11.544487

**Authors:** Rodrigo Costa-Araújo, Gustavo Rodrigues Canale, Fabiano Rodrigues de Melo, Raimundo Rodrigues da Silva, Ivan Baptista da Silva, Raony Macedo de Alencar, Luciano Ferreira da Silva, Leandro Jerusalinsky, Renata Bocorny de Azevedo, Eduardo Marques Santos Júnior, Italo Mourthé, Emil José Hernández Ruz, José de Sousa e Silva, Christian Roos, Izeni Pires Farias, Tomas Hrbek

## Abstract

The arc of deforestation, located between the southern Amazonia and the northern Cerrado of Brazil, is a top deforestation frontier worldwide. The high deforestation rates in this region are caused by human activities and threatens a rich diversity of primates that are still poorly-known and therefore investments into taxonomy and distribution research are urgent to support science-driven conservation of primates of this region. Here we present a dataset of 192 new occurrence records for 22 species of primates of the genera *Alouatta, Aotus, Ateles, Cebus, Chiropotes, Lagothrix, Leontocebus, Pithecia, Plecturocebus, Saimiri*, and *Sapajus*, collected during 10 field expeditions carried out across the arc of deforestation between 2015–2018. Based on these occurrence records we extend the ranges of *Saimiri collinsi, Sapajus apella*, and *Alouatta puruensis*, identify a potential hybridization zone between *A. puruensis* and *Alouatta discolor*, and redefine the range of *Plecturocebus moloch*. Moreover, this dataset is a useful source of information otherwise scarce for further researches on species distributions and habitat use patterns, on the effect of environmental variables on such patterns, for estimating population sizes, evaluating habitat suitability, for predicting effects of climate change, habitat loss and fragmentation on populations, and for assessing species extinction risks. The ranges of primates endemic to the arc of deforestation, with a few exceptions, are fragile hypotheses, which hampers the establishment of effective conservation efforts for species increasingly threatened on a global deforestation frontier.

## 1 Introduction

The arc of deforestation is a region located in southern Amazonia and northern Cerrado of Brazil where approximately half of the global forest conversion into anthropogenic landscapes has occurred in the past three decades (Fearnside et al., 2009; FAO, 2016; Silva-Jr. et al., 2019). The deforestation in this region is a direct consequence of the implementation of BR-230 (Transamazônica) and BR-163 (Cuiabá-Santarém) roadways in early 70’s, which prompted an exponential and disordered invasion of people from south and southeast Brazil motivated by land tenure opportunities, lack of enforcement of environmental laws, and production of beef, soy bean, and corn commodities (Laurance et al., 2002; Kirby et al., 2006; Fearnside et al., 2017).

On the other hand, the arc of deforestation harbours a rich diversity of primates that are poorly studied and have been directly and negatively affected by anthropogenic habitat reduction. To date, 52 primate species are known for this region, including a number of recently described taxa such as *Mico munduruku, Mico schneideri* (Costa-Araújo et al., 2019, 2021), and *Plecturocebus grovesi* (Boubli et al., 2019). The conservation status of such recently described species (Boubli et al., 2020; Costa-Araújo et al., 2020, 2022a) and of a few others studied beyond the species description––e.g. *Ateles marginatus, Cebus kaapori, Chiropotes satanas, Mico marcai, Plecturocebus vieirai*–*–*are rapidly escalating to threatened by extinction according to IUCN (Silva et al., 2018; Fialho et al., 2021; Port-Carvalho et al., 2021; Ravetta et al., 2021; Costa-Araújo et al., 2022b, c), including the case of *P. grovesi*, considered as one of the 25 most threatened primates worldwide (Boubli et al., 2022).

In order to contribute to fill the gaps of basic knowledge on and to support the conservation of primates from the arc of deforestation, we present here a dataset of 192 new occurrence records for 22 primate species. Although such type of data is currently scarce for primates of the arc of deforestation, it is paramount for assessing species conservation status and extinction risks, as well as for designing conservation measurements and for improving the effectiveness of existent strategies, for example, the National Action Plan for the Conservation of Amazonian Primates (ICMBio, 2017). Currently, the distribution of primates endemic to the arc of deforestation, with a few exceptions, are fragile hypotheses based on a handful of records located along the large Amazonian rivers.

## 2 Material and methods

During four years (2015–2018), 10 field expeditions were carried out across the arc of deforestation, southern Amazonia, in the States of Amazonas, Mato Grosso, Pará, and Rondônia, Brazil. All large river interfluves were surveyed for primate occurrence as well as interfluves formed by rivers of second order along the basins of the Xingu, Tapajós, Aripuanã, Ji-Paraná, Teles Pires, Juruena, Madeira, and Amazonas Rivers (Fig. 1). In each expedition (21 days in average) active search for primates and playback survey for the detection of marmosets were carried out by RCA and usually one assistant. For each primate observation or vocalization, the species was recorded and the exact geographical coordinates were taken with a Garmin GPSMap 65s device. Species identification follows the latest taxonomic review of Neotropical primates (Rylands & Mittermeier, 2013) and are based on the phenotype or on the locality, as in the case of vocalization records. All the records, in WGS 84 ellipsoid and GMS datum, were posteriorly transformed in decimal degrees using the speciesLink geographical coordinates converter (http://splink.cria.org.br/conversor?criaLANG=en) and plotted on a map using QGIS (2023).

**Figure 1.**
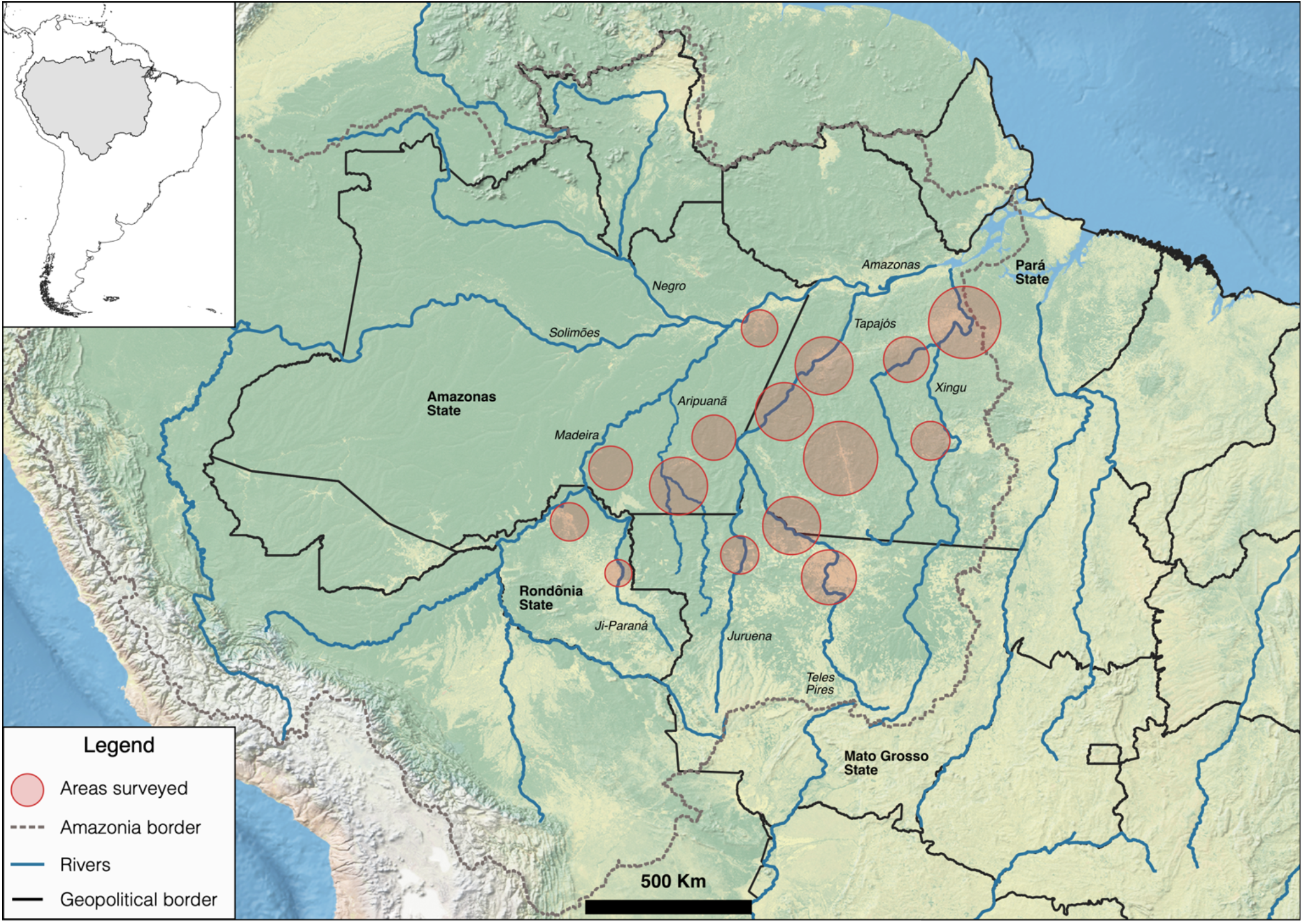
**Areas surveyed for primate occurrence during 10 field expeditions carried out between 2015–2018 in the arc of deforestation, southern Brazilian Amazonia.**

## 3 Results and discussion

The dataset presented here brings 192 new records of 22 primate species of the genera *Alouatta, Aotus, Ateles, Cebus, Chiropotes, Lagothrix, Leontocebus, Pithecia, Plecturocebus, Saimiri*, and *Sapajus* (Supplementary Material), in addition to records from the same field expeditions previously published elsewhere (Costa-Araújo et al., 2019, 2021, 2022; Boubli et al., 2019; Byrne et al., 2021) or subject of specific ongoing research. Such dataset is an useful source of information for further researches on the geographic distribution and habitat use, the effect of environmental variables on such patterns, on population size estimation and habitat suitability evaluation, as well as for predicting the effects of climate change, habitat loss and fragmentation on populations, and for assessing the risk of extinction of these 22 primate species.

Based on current knowledge on the species range limits (IUCN, 2022), our dataset sheds light into the distribution and population dynamics of five primate species of the arc of deforestation. *Saimiri collinsi* was recorded on the left bank of Jamanxim River, and on the right bank of the upper Tapajós and lower Teles Pires Rivers, extending the known species distribution for approximately 600 Km to the West (Fig. 2a). *Sapajus apella* was recorded on the upper Teles Pires River, which extends the species distribution for approximately 100 Km to the Southeast (Fig. 2b).

**Figure 2.**
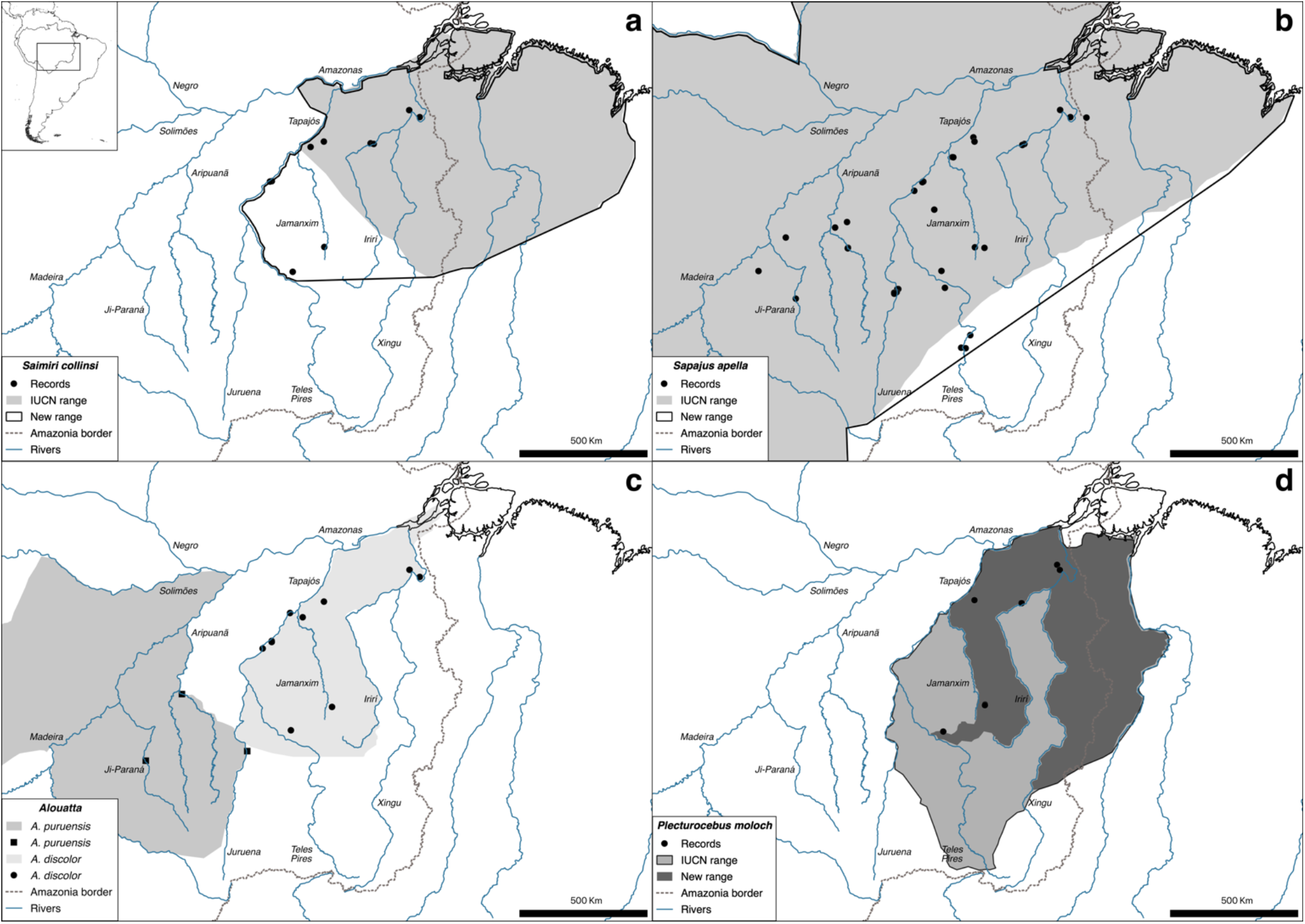
**Changes in the distribution of four primate species on the arc of deforestation, southern Amazonia, Brazil: a. 600 Km west extension of the range of *Saimiri collinsi*; b. 100 Km southeast extension of the range of *Sapajus apella*; c. extension of the range of *Alouatta puruensis* into the Juruena–Teles Pires interfluve, a probable hybridization zone with *A. discolor*; d. new provisional range proposed for *Plecturocebus moloch*.**

One of the records of *A. puruensis* (Fig. 3), on the right bank of the medium Juruena River, extends the species distribution into the Juruena–Teles Pires interfluve, a locality only 20 Km distant from the southwestern limit of the distribution of *A. discolor* (Fig. 2c). Although there is a record of sympatry between these two *Alouatta* species in Paranaíta municipality (Pinto and Setz, 2000), the only species formerly considered to occur in the Juruena–Teles Pires interfluve is *A. discolor* (see also Gregorin, 2006). We suggest that the northern portion of such interfluve is a natural hybridization zone for *A. puruensis* and *A. discolor* as introgression is relatively common in primates (Cortés-Ortiz et al., 2019a) and expected in sympatric howler monkey species (e.g. Cortés-Ortiz et al., 2019b; Mourthé et al., 2019). Further field work is needed to gather occurrence records and samples for genetic and taxonomic studies that could determine the extent of occurrence of *A. puruensis* and *A. discolor* in this region, and confirm the existence of and characterize the population dynamics on the hybrid zone.

**Figure 3.**
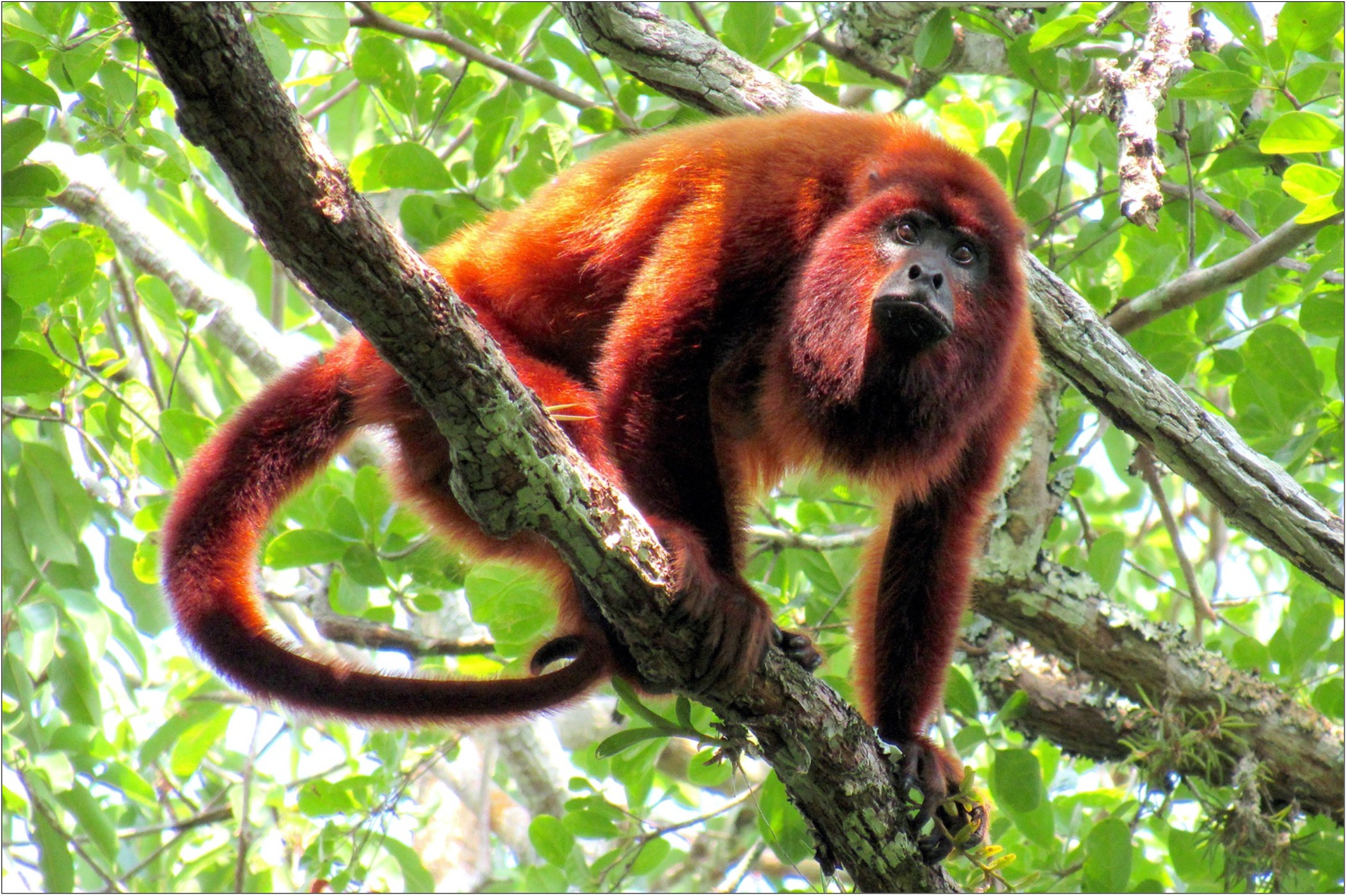
**Adult *Alouatta puruensis* recorded on the right bank of the medium Juruena River, Nova Bandeirantes, Mato Grosso State, Brazil.**

We propose a new hypothesis for the geographical distribution of *P. moloch* (Fig. 2d) based on our records for the species and subtracting the ranges of *P. grovesi* (Boubli et al., 2019) and *P. vieirai* (Costa-Araújo et al., 2022), described from titi populations within the range originally proposed for *P. moloch*. Nonetheless, our hypothesis is provisional given that the taxonomy of *P. moloch* still demands further investigation to determine the species type locality, the validity of two species synonyms, *P. remulus* and *P. emiliae*, and the taxonomy of titis from the Jamanxim–Teles Pires interfluve––this region was not included as part of the range of *P. moloch* given ongoing research (Costa-Araújo et al., in prep.) suggests they might represent a distinct species.

Finally, our record of *P. baptista* is located within the range currently hypothesized for *P. hoffmannsi*, as observed also by Rocha et al. (2019) and Printes et al. (2018). Further field surveys and taxonomic research are necessary to clarify if there is a natural hybrid zone between these two species or if these titis represent a clinal variation of a single species––as it is currently hypothesized for *P. cinerascens* and the recently described *P. parecis* (Byrne et al., 2021).

Basic knowledge on species is paramount for biodiversity protection because they are the main subjects of awareness raising and conservation strategies, and the lack of information on species taxonomy and distribution impedes effective conservation measurements (Costa et al., 2005; Rylands and Mittermeier, 2014). The implementation and improvement of conservation strategies envisioned in the National Action Plan for the Conservation of Amazonian Primates (ICMBio, 2017), for example, depend on such basic information. Therefore, given the scarcity of information on and the pace of habitat loss faced by the primates on the arc of deforestation, they should be priority for research and conservation among platyrrhines. Given that primates are valuable as flagship species (Dietz et al., 1994; Estrada et al., 2017; Chapman et al., 2020), investments on research and conservation of primates from the arc of deforestation are expected to have a positive cascade effect on the protection of Amazonian biodiversity, ecosystem services, and mitigation of climate change.

## Data availability

No datasets were used in this article.

## Supplement

The dataset of occurrence records of primates from the arc of deforestation can be accessed via the following link.

## Author contributions

Data collection was led by RCA with collaborations of TH, RRS, IBS, LFS, RMA, LJ, RBA, EMSJr, and IM; RCA, IF, TH, GRC, FRM, EH, LJ, RBA, and JSSJr contributed with resources and logistic support for the field expeditions; RCA assembled the dataset, designed the figures, and wrote the text with contributions from all authors.

## Acknowledgements

This article is a result of RCA doctoral thesis carried out at the graduate program in Ecology of the National Institute of Amazon Researches, Manaus, Brazil, with the invaluable support of Brisa Araújo and a number of people, communities and friends from southern Amazonia, especially Zagaia, Banha, Catitu, Dona Eunice, Marina and João Lutz, Natal, Seu Luís, Seu Roberto, Ozébio, Reginaldo, Canguru, Pamela Sateré-Mawé and Valdinelis, Nelson and his family, Jóia, Angelisson Tenharin, Nilcelio Djahui, Rosangela Parintintin, friends from TI Cachoeira Seca, Valdir and Jacaré, Fazenda ONF São Nicolau, the Centro Nacional de Pesquisa e Conservação de Primatas Brasileiros (ICMBio/CPB), Consórcio UHE Teles Pires, Biota Projetos e Consultoria Ambiental, and the teams of Itaituba, Apuí, and Jaru bureaus of the Instituto Chico Mendes de Conservação da Biodiversidade, without whom this research would not be possible.

