## Supplementary material for "A dataset of new occurrence records of primates from the arc of deforestation, Brazil"

Supplementary Material. Dataset of 192 new occurrence records (decimal degrees, WGS 84 ellipsoid) of 22 primate species of the *Alouatta*, *Aotus*, *Ateles*, *Cebus*, *Chiropotes*, *Lagothrix*, *Leontocebus*, *Pithecia*, *Plecturocebus*, *Saimiri*, and *Sapajus* genera from the Amazonian arc of deforestation, Brazil.

| Species | Latitude | Longitude | State | Municipality | Type of record |
|---|---|---|---|---|---|
| *Alouatta belzebul* | -3,256066 | -52,101795 | Pará | Altamira | observation |
| *Alouatta belzebul* | -3,431207 | -51,273499 | Pará | Anapu | observation |
| *Alouatta discolor* | -3,154029 | -52,236515 | Pará | Altamira | vocalization |
| *Alouatta discolor* | -3,415871 | -51,861140 | Pará | Vitória do Xingu | vocalization |
| *Alouatta discolor* | -4,321818 | -55,376763 | Pará | Rurópolis | observation |
| *Alouatta discolor* | -4,739434 | -56,620976 | Pará | Itaituba | observation |
| *Alouatta discolor* | -4,896320 | -56,157203 | Pará | Trairão | observation |
| *Alouatta discolor* | -5,766087 | -57,293432 | Pará | Jacareacanga | observation |
| *Alouatta discolor* | -5,806025 | -57,293842 | Pará | Jacareacanga | observation |
| *Alouatta discolor* | -6,043952 | -57,625627 | Pará | Jacareacanga | vocalization |
| *Alouatta discolor* | -8,188168 | -55,080342 | Pará | Altamira | observation |
| *Alouatta discolor* | -9,045870 | -56,589056 | Pará | Jacareacanga | observation |
| *Alouatta puruensis* | -7,715710 | -60,587942 | Amazonas | Novo Aripuanã | observation |
| *Alouatta puruensis* | -9,810552 | -58,192113 | Mato Grosso | Nova Bandeirantes | observation |
| *Alouatta puruensis* | -10,162898 | -61,914299 | Rondônia | Ouro Preto do Oeste | observation |
| *Aotus infulatus* | -4,381089 | -53,657286 | Pará | Altamira | observation |
| *Ateles chamek* | -6,798492 | -59,055495 | Amazonas | Apuí | observation |
| *Ateles chamek* | -6,799800 | -59,066411 | Amazonas | Apuí | observation |
| *Ateles chamek* | -7,164853 | -59,873720 | Amazonas | Apuí | observation |
| *Ateles chamek* | -7,190935 | -59,866180 | Amazonas | Apuí | observation |
| *Ateles chamek* | -7,479167 | -60,660556 | Amazonas | Apuí | observation |
| *Ateles chamek* | -7,532118 | -60,474626 | Amazonas | Apuí | observation |
| *Ateles chamek* | -7,724356 | -60,570772 | Amazonas | Apuí | observation |
| *Ateles chamek* | -7,852230 | -62,262647 | Amazonas | Humaitá | observation |
| *Ateles chamek* | -8,179956 | -60,461448 | Amazonas | Novo Aripuanã | observation |
| *Ateles chamek* | -8,232795 | -60,031846 | Amazonas | Apuí | observation |
| *Ateles chamek* | -9,655326 | -61,838057 | Rondônia | Vale do Anari | observation |

| Species | Latitude | Longitude | State | Municipality | Type of record |
|---|---|---|---|---|---|
| *Ateles chamek* | -9,683964 | -56,465441 | Mato Grosso | Paranaíta | observation |
| *Ateles chamek* | -9,718700 | -58,195878 | Mato Grosso | Nova Bandeirantes | observation |
| *Ateles chamek* | -11,420282 | -55,535212 | Mato Grosso | Sinop | observation |
| *Ateles chamek* | -11,900939 | -55,704336 | Mato Grosso | Sorriso | observation |
| *Ateles marginatus* | -4,371760 | -53,661465 | Pará | Altamira | observation |
| *Ateles marginatus* | -4,740682 | -56,619694 | Pará | Itaituba | observation |
| *Ateles marginatus* | -4,891533 | -56,158671 | Pará | Trairão | observation |
| *Ateles marginatus* | -5,649224 | -57,239382 | Pará | Itaituba | observation |
| *Ateles marginatus* | -5,764504 | -57,296657 | Pará | Jacareacanga | observation |
| *Ateles marginatus* | -6,816851 | -56,858321 | Pará | Jacareacanga | observation |
| *Ateles marginatus* | -8,089851 | -54,991485 | Pará | Altamira | observation |
| *Ateles marginatus* | -8,097535 | -54,989894 | Pará | Altamira | observation |
| *Ateles marginatus* | -8,181511 | -55,369755 | Pará | Novo Progresso | observation |
| *Ateles marginatus* | -8,191731 | -55,068541 | Pará | Altamira | observation |
| *Ateles marginatus* | -8,193925 | -55,059866 | Pará | Altamira | observation |
| *Ateles marginatus* | -8,194418 | -55,362012 | Pará | Novo Progresso | observation |
| *Ateles marginatus* | -9,059008 | -56,531378 | Pará | Jacareacanga | observation |
| *Ateles marginatus* | -9,096743 | -56,520596 | Pará | Jacareacanga | observation |
| *Ateles marginatus* | -9,104173 | -56,477179 | Pará | Jacareacanga | observation |
| *Ateles marginatus* | -11,420268 | -55,535448 | Mato Grosso | Sinop | observation |
| *Cebus unicolor* | -7,494167 | -60,482322 | Amazonas | Apuí | observation |
| *Cebus unicolor* | -7,614548 | -60,665216 | Amazonas | Apuí | observation |
| *Chiropotes albinasus* | -4,371760 | -53,661465 | Pará | Altamira | vocalization |
| *Chiropotes albinasus* | -4,377047 | -53,664946 | Pará | Altamira | observation |
| *Chiropotes albinasus* | -4,740857 | -56,619720 | Pará | Itaituba | observation |
| *Chiropotes albinasus* | -4,740935 | -56,638851 | Pará | Itaituba | observation |
| *Chiropotes albinasus* | -5,769441 | -57,291435 | Pará | Jacareacanga | observation |
| *Chiropotes albinasus* | -5,781748 | -57,261809 | Pará | Itaituba | observation |
| *Chiropotes albinasus* | -6,785360 | -59,049433 | Amazonas | Apuí | observation |
| *Chiropotes albinasus* | -6,798525 | -59,055482 | Amazonas | Apuí | observation |

| Species | Latitude | Longitude | State | Municipality | Type of record |
|---|---|---|---|---|---|
| *Chiropotes albinasus* | -7,190322 | -59,864595 | Amazonas | Apuí | observation |
| *Chiropotes albinasus* | -7,484720 | -60,477481 | Amazonas | Apuí | observation |
| *Chiropotes albinasus* | -7,501831 | -60,481858 | Amazonas | Apuí | observation |
| *Chiropotes albinasus* | -7,576732 | -60,671287 | Amazonas | Apuí | observation |
| *Chiropotes albinasus* | -7,612338 | -60,664355 | Amazonas | Apuí | observation |
| *Chiropotes albinasus* | -8,018963 | -60,136411 | Amazonas | Apuí | observation |
| *Chiropotes albinasus* | -8,125647 | -54,998307 | Pará | Altamira | observation |
| *Chiropotes albinasus* | -8,183114 | -55,365275 | Pará | Novo Progresso | observation |
| *Chiropotes albinasus* | -8,231920 | -60,029423 | Amazonas | Apuí | observation |
| *Chiropotes albinasus* | -8,232070 | -60,033680 | Amazonas | Apuí | observation |
| *Chiropotes albinasus* | -9,096868 | -56,520560 | Pará | Jacareacanga | observation |
| *Chiropotes albinasus* | -9,845558 | -58,235430 | Mato Grosso | Cotriguaçu | observation |
| *Chiropotes albinasus* | -9,911144 | -61,942478 | Rondônia | Vale do Anari | observation |
| *Lagothrix cana* | -2,956981 | -58,014683 | Amazonas | Urucurituba | observation |
| *Lagothrix cana* | -6,531828 | -59,089128 | Amazonas | Apuí | observation |
| *Lagothrix cana* | -7,163810 | -59,871508 | Amazonas | Apuí | observation |
| *Lagothrix cana* | -7,194129 | -59,864045 | Amazonas | Apuí | observation |
| *Lagothrix cana* | -7,529642 | -60,475932 | Amazonas | Apuí | observation |
| *Lagothrix cana* | -7,612338 | -60,664355 | Amazonas | Apuí | observation |
| *Lagothrix cana* | -7,736875 | -60,510270 | Amazonas | Apuí | observation |
| *Lagothrix cana* | -7,753777 | -60,366578 | Amazonas | Apuí | observation |
| *Lagothrix cana* | -7,773209 | -60,564321 | Amazonas | Apuí | observation |
| *Lagothrix cana* | -7,845560 | -62,306463 | Amazonas | Humaitá | observation |
| *Lagothrix cana* | -9,557020 | -61,604184 | Rondônia | Vale do Anari | observation |
| *Lagothrix cana* | -10,070073 | -61,969437 | Rondônia | Ji-Paraná | observation |
| *Lagothrix cana* | -10,122558 | -61,902792 | Rondônia | Ji-Paraná | vocalization |
| *Leontocebus weddelli* | -8,769543 | -63,475592 | Rondônia | Candeias do Jamari | observation |
| *Leontocebus weddelli* | -9,066980 | -63,304194 | Rondônia | Itapuã do Oeste | observation |
| *Pithecia mittermeieri* | -7,262815 | -60,061120 | Amazonas | Apuí | observation |
| *Pithecia mittermeieri* | -7,463472 | -62,932391 | Amazonas | Humaitá | observation |

| Species | Latitude | Longitude | State | Municipality | Type of record |
|---|---|---|---|---|---|
| *Pithecia mittermeieri* | -7,530242 | -60,475596 | Amazonas | Apuí | observation |
| *Pithecia mittermeieri* | -7,736256 | -60,515709 | Amazonas | Apuí | observation |
| *Pithecia mittermeieri* | -8,232976 | -60,030847 | Amazonas | Apuí | observation |
| *Pithecia mittermeieri* | -10,050181 | -58,383100 | Mato Grosso | Cotriguaçu | observation |
| *Plecturocebus baptista* | -3,422861 | -57,685694 | Amazonas | Maués | observation |
| *Plecturocebus bernhardi* | -7,456201 | -62,939055 | Amazonas | Humaitá | observation |
| *Plecturocebus bernhardi* | -7,462714 | -62,932357 | Amazonas | Humaitá | observation |
| *Plecturocebus bernhardi* | -7,838514 | -62,293781 | Amazonas | Humaitá | observation |
| *Plecturocebus bernhardi* | -7,859012 | -62,258619 | Amazonas | Humaitá | observation |
| *Plecturocebus bernhardi* | -10,063351 | -61,960871 | Rondônia | Ji-Paraná | observation |
| *Plecturocebus bernhardi* | -10,124698 | -61,908830 | Rondônia | Ji-Paraná | observation |
| *Plecturocebus bernhardi* | -10,144905 | -61,907793 | Rondônia | Ji-Paraná | observation |
| *Plecturocebus brunneus* | -8,769542 | -63,475592 | Rondônia | Candeias do Jamari | observation |
| *Plecturocebus brunneus* | -8,769543 | -63,475592 | Rondônia | Candeias do Jamari | observation |
| *Plecturocebus brunneus* | -9,066980 | -63,304194 | Rondônia | Itapuã do Oeste | observation |
| *Plecturocebus cinerascens* | -6,799215 | -59,055320 | Amazonas | Apuí | observation |
| *Plecturocebus cinerascens* | -7,204571 | -60,018139 | Amazonas | Apuí | observation |
| *Plecturocebus cinerascens* | -7,519522 | -60,653148 | Amazonas | Apuí | observation |
| *Plecturocebus cinerascens* | -7,526585 | -60,653049 | Amazonas | Apuí | vocalization |
| *Plecturocebus cinerascens* | -9,911395 | -58,377819 | Mato Grosso | Cotriguaçu | observation |
| *Plecturocebus grovesi* | -9,667538 | -56,496601 | Mato Grosso | Paranaíta | observation |
| *Plecturocebus grovesi* | -9,791949 | -58,195175 | Mato Grosso | Nova Bandeirantes | vocalization |
| *Plecturocebus hoffmannsii* | -5,741886 | -57,361631 | Pará | Jacareacanga | observation |
| *Plecturocebus hoffmannsii* | -4,942477 | -56,770202 | Pará | Itaituba | observation |
| *Plecturocebus miltoni* | -7,795544 | -60,537823 | Amazonas | Apuí | observation |
| *Plecturocebus miltoni* | -8,172043 | -60,453659 | Amazonas | Novo Aripuanã | observation |
| *Plecturocebus miltoni* | -8,176226 | -60,450408 | Amazonas | Apuí | observation |
| *Plecturocebus miltoni* | -8,178109 | -60,455417 | Amazonas | Novo Aripuanã | observation |
| *Plecturocebus miltoni* | -8,220575 | -60,025239 | Amazonas | Apuí | observation |
| *Plecturocebus miltoni* | -8,226501 | -60,029947 | Amazonas | Apuí | observation |

| Species | Latitude | Longitude | State | Municipality | Type of record |
|---|---|---|---|---|---|
| *Plecturocebus moloch* | -2,969967 | -52,349933 | Pará | Altamira | observation |
| *Plecturocebus moloch* | -3,157132 | -52,242953 | Pará | Altamira | observation |
| *Plecturocebus moloch* | -3,159748 | -52,248455 | Pará | Altamira | observation |
| *Plecturocebus moloch* | -4,268417 | -55,382044 | Pará | Rurópolis | observation |
| *Plecturocebus moloch* | -4,268418 | -55,382045 | Pará | Rurópolis | observation |
| *Plecturocebus moloch* | -4,368058 | -53,649794 | Pará | Altamira | observation |
| *Plecturocebus moloch* | -4,370280 | -53,658815 | Pará | Altamira | observation |
| *Plecturocebus moloch* | -4,372368 | -53,662517 | Pará | Altamira | observation |
| *Plecturocebus moloch* | -4,376896 | -53,649244 | Pará | Altamira | observation |
| *Plecturocebus moloch* | -8,098461 | -54,994222 | Pará | Altamira | observation |
| *Plecturocebus moloch* | -9,094306 | -56,527142 | Pará | Jacareacanga | observation |
| *Plecturocebus moloch* | -8,098461 | -54,994223 | Pará | Altamira | observation |
| *Plecturocebus moloch* | -8,125651 | -54,998562 | Pará | Altamira | observation |
| *Saimiri collinsi* | -3,159748 | -52,248455 | Pará | Altamira | observation |
| *Saimiri collinsi* | -3,414968 | -51,862088 | Pará | Vitória do Xingu | observation |
| *Saimiri collinsi* | -4,313622 | -55,379458 | Pará | Rurópolis | observation |
| *Saimiri collinsi* | -4,377047 | -53,664946 | Pará | Altamira | observation |
| *Saimiri collinsi* | -4,400933 | -53,548912 | Pará | Altamira | observation |
| *Saimiri collinsi* | -4,512367 | -55,866501 | Pará | Itaituba | observation |
| *Saimiri collinsi* | -5,767912 | -57,290098 | Pará | Jacareacanga | observation |
| *Saimiri collinsi* | -5,781224 | -57,358087 | Pará | Jacareacanga | observation |
| *Saimiri collinsi* | -8,183433 | -55,363710 | Pará | Novo Progresso | observation |
| *Saimiri collinsi* | -9,095654 | -56,522555 | Pará | Jacareacanga | observation |
| *Saimiri ustus* | -2,952100 | -58,068181 | Amazonas | Urucurituba | observation |
| *Saimiri ustus* | -6,573147 | -59,081904 | Amazonas | Apuí | observation |
| *Saimiri ustus* | -6,687303 | -59,080460 | Amazonas | Apuí | observation |
| *Saimiri ustus* | -7,722717 | -60,574936 | Amazonas | Apuí | observation |
| *Saimiri ustus* | -7,856605 | -62,262775 | Amazonas | Humaitá | observation |
| *Saimiri ustus* | -8,227582 | -60,032353 | Amazonas | Apuí | observation |
| *Saimiri ustus* | -8,839395 | -63,937128 | Rondônia | Porto Velho | observation |

| Species | Latitude | Longitude | State | Municipality | Type of record |
|---|---|---|---|---|---|
| *Saimiri ustus* | -9,067489 | -63,304714 | Rondônia | Itapuã do Oeste | observation |
| *Saimiri ustus* | -9,844137 | -58,233728 | Mato Grosso | Cotriguaçu | observation |
| *Saimiri ustus* | -9,848728 | -58,269594 | Mato Grosso | Cotriguaçu | observation |
| *Saimiri ustus* | -9,863258 | -58,257559 | Mato Grosso | Cotriguaçu | observation |
| *Saimiri ustus* | -9,868031 | -58,328950 | Mato Grosso | Cotriguaçu | observation |
| *Sapajus apella* | -3,153210 | -52,238517 | Pará | Altamira | observation |
| *Sapajus apella* | -3,414968 | -51,862088 | Pará | Vitória do Xingu | observation |
| *Sapajus apella* | -3,433550 | -51,272531 | Pará | Anapu | observation |
| *Sapajus apella* | -4,158472 | -55,420808 | Pará | Rurópolis | observation |
| *Sapajus apella* | -4,311525 | -55,379610 | Pará | Rurópolis | observation |
| *Sapajus apella* | -4,313622 | -55,379458 | Pará | Rurópolis | observation |
| *Sapajus apella* | -4,400933 | -53,548912 | Pará | Altamira | observation |
| *Sapajus apella* | -4,435548 | -53,628088 | Pará | Altamira | observation |
| *Sapajus apella* | -4,438924 | -53,621913 | Pará | Altamira | observation |
| *Sapajus apella* | -4,439817 | -53,623675 | Pará | Altamira | observation |
| *Sapajus apella* | -4,884648 | -56,187969 | Pará | Trairão | observation |
| *Sapajus apella* | -4,890649 | -56,156454 | Pará | Trairão | observation |
| *Sapajus apella* | -5,781852 | -57,268322 | Pará | Itaituba | observation |
| *Sapajus apella* | -5,783523 | -57,256974 | Pará | Itaituba | observation |
| *Sapajus apella* | -5,805452 | -57,292603 | Pará | Jacareacanga | observation |
| *Sapajus apella* | -6,121724 | -57,589114 | Pará | Jacareacanga | observation |
| *Sapajus apella* | -6,812682 | -56,855636 | Pará | Jacareacanga | observation |
| *Sapajus apella* | -7,268159 | -60,057074 | Amazonas | Apuí | observation |
| *Sapajus apella* | -7,470663 | -60,494342 | Amazonas | Apuí | observation |
| *Sapajus apella* | -7,836196 | -62,303610 | Amazonas | Humaitá | observation |
| *Sapajus apella* | -8,195322 | -55,366323 | Pará | Novo Progresso | observation |
| *Sapajus apella* | -8,211416 | -60,020714 | Amazonas | Apuí | observation |
| *Sapajus apella* | -8,215703 | -55,014563 | Pará | Altamira | observation |
| *Sapajus apella* | -8,227582 | -60,032353 | Amazonas | Apuí | observation |
| *Sapajus apella* | -8,232947 | -60,029154 | Amazonas | Apuí | observation |

| Species | Latitude | Longitude | State | Municipality | Type of record |
|---|---|---|---|---|---|
| *Sapajus apella* | -9,063070 | -56,592167 | Pará | Jacareacanga | vocalization |
| *Sapajus apella* | -9,067489 | -63,304714 | Rondônia | Itapuã do Oeste | observation |
| *Sapajus apella* | -9,682842 | -56,465968 | Mato Grosso | Paranaíta | observation |
| *Sapajus apella* | -9,727820 | -58,175349 | Mato Grosso | Nova Bandeirantes | observation |
| *Sapajus apella* | -9,730757 | -58,210872 | Mato Grosso | Nova Bandeirantes | observation |
| *Sapajus apella* | -9,846406 | -58,236330 | Mato Grosso | Cotriguaçu | observation |
| *Sapajus apella* | -9,848728 | -58,269594 | Mato Grosso | Cotriguaçu | observation |
| *Sapajus apella* | -9,856182 | -58,260230 | Mato Grosso | Cotriguaçu | observation |
| *Sapajus apella* | -9,863445 | -58,328272 | Mato Grosso | Cotriguaçu | observation |
| *Sapajus apella* | -9,898467 | -58,323943 | Mato Grosso | Cotriguaçu | observation |
| *Sapajus apella* | -10,080955 | -61,937099 | Rondônia | Ji-Paraná | observation |
| *Sapajus apella* | -11,420268 | -55,535448 | Mato Grosso | Sinop | observation |
| *Sapajus apella* | -11,420282 | -55,535212 | Mato Grosso | Sinop | observation |
| *Sapajus apella* | -11,881577 | -55,868323 | Mato Grosso | Sorriso | observation |
| *Sapajus apella* | -11,900939 | -55,704336 | Mato Grosso | Sorriso | observation |
